# The Impact of Brain-Derived Neurotrophic Factor rs6265 (Val66Met) Polymorphism on Therapeutic Electrical Stimulation for Peripheral Nerve Regeneration: A Preclinical Study of Therapy-Genotype Interactions

**DOI:** 10.1101/2024.08.29.610333

**Authors:** Jordan Walters, Maria J. Quezada, Suning He, Kathy Steece-Collier, Timothy J. Collier, Caryl E. Sortwell, Colin K. Franz

**Author notes:** Correspondence: Colin K. Franz, MD, PhD, FAAPMR Physician-Scientist, Shirley Ryan AbilityLab Associate Professor of Physical Medicine and Rehabilitation, and Neurology, Northwestern University 355 E. Erie St, Chicago, IL 60611.

## Abstract

The Impact of Brain-Derived Neurotrophic Factor rs6265 (Val66Met) Polymorphism on Therapeutic Electrical Stimulation for Peripheral Nerve Regeneration: A Preclinical Study of Therapy-Genotype Interactions

**Introduction:** Peripheral nerve injuries (PNIs) significantly impact patient quality of life. Therapeutic electrical stimulation (TES) shows promise in enhancing nerve regeneration, but outcomes vary widely. This study investigates the impact of the rs6265 single nucleotide polymorphism (SNP) on TES efficacy in a preclinical rat model and human stem cell-derived motor neurons.

**Methods:** Wild-type (WT) and rs6265 variant rats underwent sciatic nerve transection and received either TES or sham treatment. Muscle reinnervation was assessed through compound muscle action potentials and muscle fiber cross-sectional area. Isogenic human iPSC-derived motor neurons were used to study activity-dependent BDNF secretion.

**Results:** TES improved muscle reinnervation and fiber size in WT but not rs6265 allele carriers. rs6265 allele carriers exhibited impaired activity-dependent BDNF secretion in vitro.

**Discussion:** The rs6265 polymorphism influences TES efficacy, highlighting the need for personalized approaches in PNI treatment. These findings suggest that genetic screening could optimize therapeutic outcomes.

**Clinical Relevance:** Understanding genetic factors affecting TES response can enhance treatment strategies for PNI, potentially improving patient recovery and reducing outcome variability.

## Introduction

Peripheral Nerve Injuries (PNIs) are characterized by a significant detriment to patients’ quality of life and represent a formidable challenge in neuromuscular medicine, affecting an estimated 200,000 individuals annually in the United States^1,2^. Despite some capacity for peripheral axons to regenerate, the majority of people affected by PNI experience slow and incomplete functional recovery. Incomplete recovery can result in permanent disability, severe neuropathic pain, and necessitate invasive surgeries like free functioning muscle grafts to salvage modest limb function^3^. The inherently slow pace of axon regeneration represents the greatest challenge, which is inadequate when faced with the large clinical distances required for effective healing, including nerve injuries localized to the proximal limb^4^. This situation is exacerbated by the absence of any FDA-approved pharmacological interventions to enhance axon regrowth, underscoring a dire need for innovative therapeutic strategies.

Therapeutic electrical stimulation (TES) has emerged as a promising intervention to promote axon regrowth and improve muscle reinnervation. Generally characterized by a single session of low-frequency stimulation, TES was developed in a rat model of PNI^5,6^ and has shown promise in several small human randomized control trials^7–10^. However, the variability in clinical outcomes has been a significant hurdle, likely influenced by factors such as age, sex, and medical comorbidities such as diabetes mellitus^11,12^. Yet, the interplay between patient genetic variations and PNI recovery remains a largely uncharted territory. PNI clinical studies lack the scale necessary for genome-wide association studies, which require large sample sizes to detect genetic variations associated with treatment responses^13^. This limitation has steered research towards a proof-of-concept approach in a preclinical model, focusing on specific genes of interest rather than broad genomic analysis. To this end, a particularly intriguing genetic candidate is the rs6265 single nucleotide polymorphism (SNP) in the brain-derived neurotrophic factor (BDNF) gene, also known as Val66Met in humans. This genetic variant has a high, but geographically heterogeneous, global prevalence of more > 20% (e.g., 22% in Caucasian populations, 72% Asian, 2% African American)^14–17^. The rs6265 SNP is understood to impair BDNF secretion in response to neural activity, which is implicated as a central mechanism in the effect of TES on axon regeneration^18,19^. Additionally, since BDNF plays a crucial role in recovery after PNI, there is a theoretical risk posed by rs6265 on the efficacy of TES^20^. The high frequency of this genetic trait in the population enforces the importance of defining a gene-treatment interaction, as well as the clinical relevance of this issue.

Our study aims to elucidate the effects of rs6265 on the efficacy of TES in promoting nerve regeneration in preclinical models. Here, we mainly focus on a rat model harboring the rs6265 SNP, which undergoes sciatic nerve transection and repair. It is important to note that rs6265 is known as Val66Met in humans, but Val68Met in rodents. The difference in numbering arises due to slight variations in the amino acid sequence near the polymorphism site between the two species^21^. To isolate the impaired ability of rs6265 motor neurons to secrete BDNF, we conducted experiments in a human pluripotent stem cell-derived model of rs6265, where we demonstrated that the rs6265 motor neurons have impaired activity-dependent BDNF release. By focusing on this specific gene, we seek to demonstrate proof-of-concept on how genetic factors can influence clinically inspired treatment outcomes for PNI. This investigation represents a vital step towards developing more precise and effective treatments for PNI with the potential of reducing outcome variability and enhancing recovery for those affected.

## Methods

All animal use procedures were approved by Northwestern University’s Institutional Animal Care and Use Committee (IACUC) and performed in full compliance with the National Institutes of Health Guide for the Care and Use of Laboratory Animals and the guidelines of the National Society for Medical Research. Rats were housed in Northwestern University’s animal care facilities on a 12/12-hour light cycle, with food and water available ad libitum. Pain and distress were monitored and controlled appropriately by study personnel and veterinary staff.

## Treatments

Wild-type (WT; n = 25) and rs6265 (n = 25) rats of both sexes underwent transection and direct repair of the sciatic nerve. The CRISPR knock-in rat model of the rs6265 BDNF SNP used in this study has been previously described in detail^21^. Rats were then subjected to either one hour of TES (20 Hz, 3V) proximal to the injury site (n = 23) or a sham treatment (n = 27). Sham animals received no electrical stimulation while under anesthesia for one hour, after nerve injury and before wound closure.

## Surgery

During surgery, rats were anesthetized using isoflurane gas (3% for induction and 1-2% for maintenance), given preoperative analgesia (SQ: Meloxicam, 1 mg/kg), and placed on a warming pad for body temperature maintenance. The sciatic nerve was exposed and transected, then directly repaired end-to-end with 10-0 nylon suture. After injury, animals received either TES proximal to the transection or a sham treatment. After one hour, the muscle incision was stitched using 5-0 absorbable sutures, and the skin incision was closed with wound clips (Autoclip system, Fine Science Tools).

## Therapeutic Electrical Stimulation

Immediately following sciatic nerve transection and repair, one hour of continuous 20 Hz TES with 200 µs pulse width was delivered at 3V, powered and controlled by a Grass SD9 Stimulator (Grass Instruments Co., Quincy, MA).

## Compound Muscle Action Potentials

Measurements of evoked compound muscle action potential (CMAP) amplitude were made with a 1”, 30 G concentric needle electrode connected to a clinical grade Nicolet EDX system with Synergy software (Natus Neurology, Middleton, WI). Animals were anesthetized during the procedure, and CMAP amplitude measurements of medial gastrocnemius (MG) and lateral gastrocnemius muscles (LGs) were taken at baseline, four-, and eight-weeks.

## Ultrasound Cross-Sectional Area

Ultrasound images were captured using a Logiq Ultrasound System (P9 XDclear, General Electric) with a Matrix Linear Array Probe (ML6-15L, General Electric). The ipsilateral cross-sectional area (CSA) was normalized to the contralateral to account for differences in animal size. Animals were anesthetized and three images of each LG were taken and averaged to attain the CSA at each of two time points (four- and eight-weeks post-transection).

## Tissue Handling

Eight weeks after surgery and treatment, animals were sacrificed and intracardially perfused with phosphate-buffered saline followed by 4% paraformaldehyde. Ipsilateral MGs and LGs were harvested and immersed in 4% paraformaldehyde for 24 hours. Then, muscles were stored in 30% sucrose solution for 24 hours, and snap-frozen in Tissue Freezing Material (TFM-C, Thermo Fisher Scientific). MGs were sectioned into 15 µm thick transverse sections on a cryostat (CM3050S, Leica) at −25 °C. LG muscles were transversely sectioned 25 µm thick on the same machine.

## Muscle Fiber Analysis

To determine the fiber cross-sectional area (FCSA), MGs were stained with Collagen IV (ab6586, Abcam) and imaged at 10x (DM6000 B, Leica). For each animal, four random sections of the muscle were captured, and 400 muscle fibers were averaged using ImageJ.

## Isogenic Human Stem Cell Model of rs6265

To isolate the effects of rs6265 on activity dependent-BDNF release by motor neurons, we employed an isogenic stem cell model. This model used stem cell lines with identical genetic backgrounds so we could study the effects of a specific genetic variant introduced through gene editing techniques. A healthy control human induced pluripotent stem cell (iPSC) line, BJFF.6, and a gene edited clone were obtained from Washington University Genome Engineering and iPSC Center (GEiC)^22^. Next generation sequencing (NGS) was used to determine that the BJFF.6 line was heterozygous for the desired rs6265 SNP in the BDNF gene. The mutation was then corrected at nucleotide 196 (G/A) using the CRISPR/Cas9 system (Figure 4A). Approximately 1 × 10^6 single cells were resuspended in P3 primary buffer (Lonza) along with a gRNA/Cas9 ribo-nuclease protein complex and the SNP correction, single-stranded oligodeoxynucleotide (ssODN). The ssODN was designed to produce the amino acid substitution from methionine to valine at codon 66. The cell suspension was electroporated. Following nucleofection, editing efficiency was assessed by NGS using primer sets specific to the target region. The cell pool was then subjected to single-cell sorting to isolate individual mutant clones. These single-cell clones were further screened with NGS analysis to confirm the status of rs6265 (Figure 4B). Next, karyotyping of the parent iPSC line and clones was completed to rule out any acquired chromosomal abnormalities using the Giemsa-Trypsin-Wright banding method. All iPSC lines were cultured in mTeSR Plus medium (STEMCELL Technology) on Matrigel-coated plates at 37°C in a humidified, 5% CO2 incubator. Cells were passaged at a 1:6 ratio using ReLeSR™ (STEMCELL Technology) to maintain their pluripotent state and ensure optimal growth conditions.

## Immunocytochemistry Staining of Isogenic iPSCs

The cultured isogenic iPSCs were stained in accordance with the Invitrogen Pluripotent Stem Cell Immunocytochemistry Kit (Thermofisher Scientific, A25526). Briefly, the culture medium was aspirated from each well of a six-well tissue culture plate and replaced with two 15-minute room temperature incubations of fixative and permeabilization solution, respectively.

Next, the iPSCs underwent a 30-minute room temperature incubation of blocking solution. Once that time elapsed, SSEA4 and OCT4 primary antibodies were diluted within the blocking solution and incubated for three hours at 4°C to show the pluripotency of the cells (figure 4C).

The cells were then washed and counterstained with secondary antibodies for one hour at room temperature. Following the final incubation, the iPSCs were washed once again, with two drops of NucBlue Fixed Cell Stain (DAPI) added to the last wash. Imaging was performed on a Nikon Crest X-Light confocal microscope (Nikon Instruments Inc.).

## Motor Neuron Differentiation

Motor neuron differentiation was conducted following the previously reported method by Ziller et al. with minor modifications^23,24^. iPSC colonies were dissociated using ReLeSR (5872; Stemcell Technologies, Vancouver, Canada) and plated at a density of 74,000 cells/cm2 in mTeSR Plus (5825; Stemcell Technologies, Vancouver, Canada) containing 10 μM ROCK inhibitor (RI, 129830-38-2, DNSK International, Hamden, CT, USA) for 24 hours.

Differentiation was initiated once cells reached 80-90% confluency by changing the base medium (N2B27 medium) to one containing 47.5 vol % DMEM:F12, 47.5 vol % Neurobasal, 1 vol % NEAA, 1 vol % Glutamax, 1 vol % N2 and 2 vol % B27. From day 0 to day 5, cells were fed daily using the N2B27 medium supplemented with 10 μM SB431542 (DNSK International), 100 nM LDN193189 (DNSK International), 1 μM Retinoic Acid (RA, Sigma-Aldrich) and 1μM of Smoothened-Agonist (SAG, DNSK International). Starting on day 6, the N2B27 culture medium supplements were changed to 1 μM RA, 1 μM SAG, 5 μM DAPT (DNSK International) and 4 μM SU5402 (DNSK International). Cells were fed daily until day 14 of differentiation.

The resulting differentiated motor neurons were then enzymatically dissociated using TrypLE Express (GIBCO, Life Technologies) supplemented with DNase I (Worthington), and further dissociated to single cells with Accutase (Sigma-Aldrich). Cells were frozen before use in our experiments. To thaw frozen motor neurons, vials were partially submerged in 37°C water bath until small pellet remained. Cells were resuspended in 4 mL of N2B27 culture medium supplemented with 10 μM RI and 2% fetal bovine serum. Then, cells were spun at 1000 rpm for 5 minutes, followed by media aspirations and cell resuspension in 1 mL of same media. Cells were counted with an automated cell counter and plated at desired densities.

## Mouse Glia Isolation

Pups were euthanized with isoflurane in a hood. After decapitation, heads and cortices were placed on ice for dissection. Skin and skull were carefully removed, followed by isolation of two cortices and striatum. Then, hippocampus, olfactory bulb, and meninges were removed from each cortex and placed in 24 well plates. Tissues were incubated at 37°C with 500 mL of pre-warm trypsin-EDTA 0.25% solution for 8 minutes, followed by tissue trituration by pipetting tissue. Glia media was prepared with 86% vol MEM, 3 vol% 20% glucose solution, 1 vol% pen/strep and 10 vol% heat inactivated horse serum and used for further glia cell isolation and expansion. To quench trypsin, 1.5 mL of glia media was added to each well, followed by further trituration with additional glia media. Suspension was spun down at 1000 rpm for 5 minutes and resuspended in 1 mL of glia media. The trituration procedure was repeated and filtered using 100 μm filter. Filtrate was then plated onto poly-D-lysine (PDL) coated 10 cm flasks at a density of 2 million cells per flask. Cells were kept in culture and fed weekly, and then further frozen or used for experiments.

## Electrophysiological Characterization of iPSC-derived motor neurons

CytoView MEA 96-well plates (Axion Biosystems) were dot coated with 20 μL of 0.1 mg/mL PDL overnight at 4°C, followed by triple PBS washes. Plates were then coated with 20 μL of 20 μg/mL laminin for 24 hours at 37°C followed by triple PBS washes. Analyzed data include four differentiation replicates for the WT (Val/Val) genotype and three differentiations for the rs6265 (Val/Met) genotype. For each differentiation replicate, 20,000 cells per well were plated in an average of 16 wells for each genotype condition. Recordings were acquired every five days post-plating using MaestroPro and AxIS Navigator (Axion BioSystems), at 37°C, 5% CO2 concentration, and using 4 kHz Kaiser Window digital low pass filter and 200 Hz IIR digital high pass filter and analyzed using AxIS Metric plotting Tool (Axion BioSystems). The spike detection method used for firing rate quantification was the adaptive threshold crossing with 6 standard deviations, with events detected only with threshold crossing between 0.84 ms and 2.26 ms pre- and post-spike, respectively. For bursting frequency characterization, an inter-spike interval threshold of 100 ms and a minimum number of 5 spikes was used. Depiction of iPSCs on multielectrode arrays is shown in Figure 4D.

## BDNF Secretion Quantification of Isogenic Human Stem Cell Model of rs6265

To measure endogenous BDNF secretion levels of isogenic lines, a total BDNF Quantikine ELISA kit (R&D Systems) was used. 96 well silicone membrane plates were treated with PDL and laminin as previously mentioned, and 5,000 glia cells and 30,000 isogenic motor neurons were then plated. 24 hours after application of either sodium channel blocker Tetrodotoxin (TTX) (10 uM, Abcam) or PBS (Sham), 50 μL of cell supernatant were collected for analysis. Optical density of each well was measured using Synergy HTX plate reader (BioTek Instruments Inc.) set at 450 nm, with wavelength correction set to 540 nm.

## Statistical Analysis

A priori power analysis (performed in G-Power) determined the number of animals needed (n=51) to adequately power a two-way ANOVA comparison with an effect size of 0.5, α = 0.05. For statistical analyses, α level was set to 0.20. Results are expressed as mean values with one standard deviation. Two-way ANOVAs were used to determine the effects of treatment and genotype. Three-way ANOVAS showed no effect of sex for each in vivo outcome measure, so data for male and female rats were combined for each treatment. Similar to the human population, our breeding strategy produced offspring with both one and two copies of the rs6265 allele. We performed a subgroup analysis using the four-week CSA data, which indicated no difference between rats with the rs6265 allele [p = 0.2515], so we grouped all rs6265 carriers together for analysis.

## Results

### Ultrasound Cross-Sectional Area: Electrical Stimulation Treatment Leads to Greater Muscle Cross-Sectional Area in Wild-Type but not rs6265 Allele Carriers

We measured the CSA of LGs four and eight weeks after nerve transection and repair. A depiction of the ultrasound image acquisition process and an example ultrasound image are shown in Figure 1A. To account for differences in body size due to sex and genotype, measurements were normalized to the contralateral limb. These normalized CSA values were compared between male and female rats at four and eight weeks after injury using a three-way ANOVA. No effect of sex was found at either time point, so male and female data were combined [p = 0.1675 for week four, p = 0.1723 for week eight] for both. A two-way ANOVA was used to determine that at four weeks, WT Estim (0.762 ± 0.046, n = 12) rats had larger LG CSA than WT Sham (0.718 ± 0.060, n = 13) [p = 0.0468]. rs6265 allele carriers had similar CSA averages regardless of treatment, as rs6265 Estim averaged 0.723 ± 0.044 (n = 11) and rs6265 Sham had a LG CSA of 0.720 ± 0.061 (n = 10) [p = 0.8117]. The same trend was evident at eight weeks, as WT Estim (0.859 ± 0.049, n = 12) rats had significantly higher values on average than WT Sham (0.788 ± 0.068, n = 13) [p = 0.0049]. Additionally, WT Estim outperformed both rs6265 Estim (0.807 ± 0.054, n = 11) and rs6265 Sham (0.808 ± 0.0644, n = 14) [p = 0.0424 for rs6265 Estim, p = 0.0341 rs6265 Sham]. rs6265 Estim and rs6265 Sham animals did not significantly differ [p = 0.9754]. Together, these indicate that electrical stimulation treatment enhances LG CSA in WT but not in rs6265 allele carriers (Figure 1B, Figure 1C).

**Figure 1:**
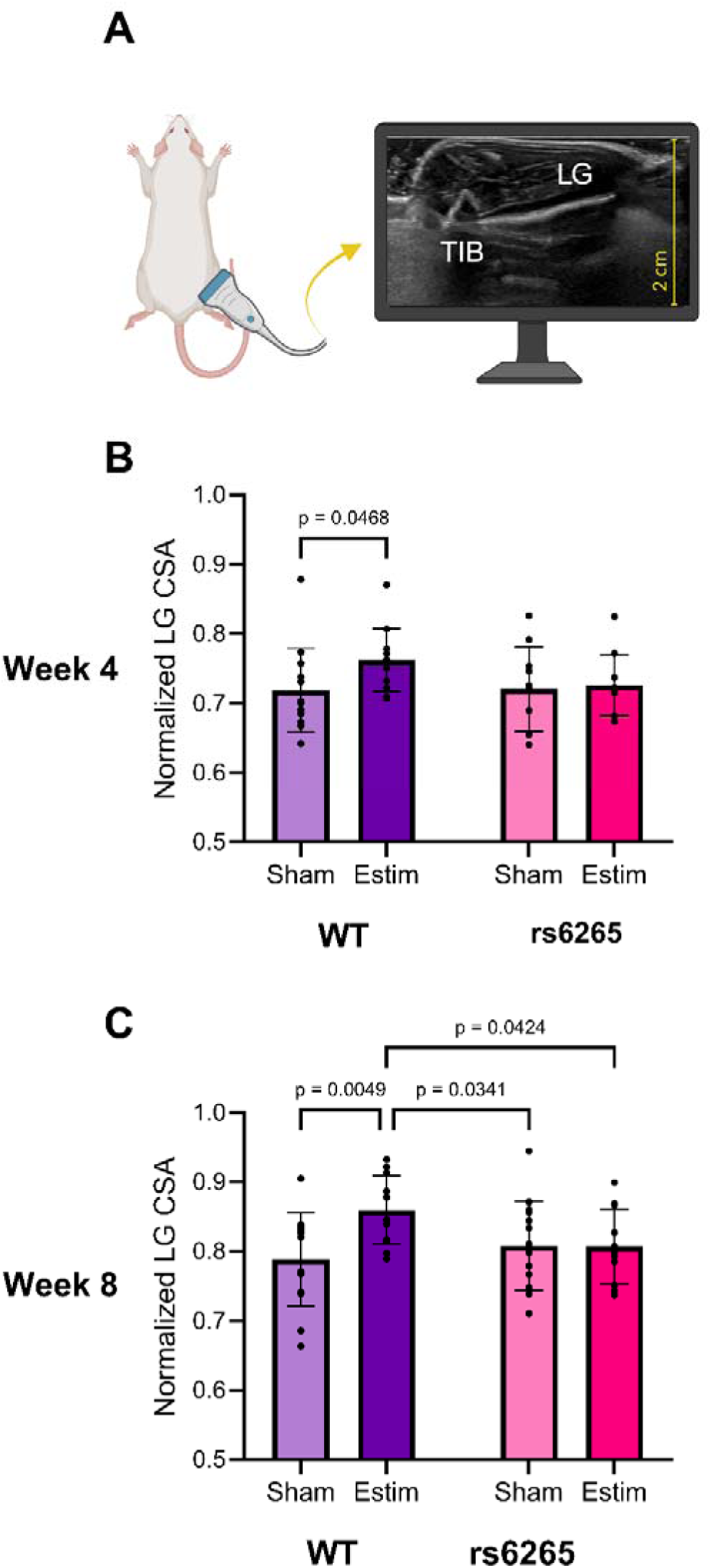
Ultrasound Cross-Sectional Area of Lateral Gastrocnemius Muscle. **A.** Depiction of ultrasound image process and example ultrasound image of lateral gastrocnemius muscle (LG). TIB= Tibia. Created with BioRender.com. **B.** Four-week ultrasound cross-sectional area (CSA) of ipsilateral LG normalized to contralateral muscle. **C.** Eight-week LG CSA normalized to contralateral.

### Compound Muscle Action Potentials: Electrical Stimulation Treatment Leads to Greater Muscle Reinnervation in Wild-Type but not rs6265 Allele Carriers

To evaluate muscle reinnervation, we evoked direct electromyographic (EMG) responses in response to sciatic nerve stimulation from the ipsilateral MGs and LGs at baseline and eight weeks post-injury. Example rectified CMAP responses recorded at that time from WT and rs6265 rodents in both Estim and Sham treatment groups are shown in Figure 2A. The rectified amplitudes of CMAP responses in all rats were normalized to the baseline value acquired before injury. Differences in these normalized CMAP amplitudes were compared between male and female rats in all four treatment groups using a three-way ANOVA. There was no effect of sex found in either muscle [p = 0.0848 for MG, p = 0.1336 for LG] so data were combined for each treatment. Using a two-way ANOVA, it was determined that WT Estim (0.591 ± 0.244, n = 12) amplitudes averaged higher than WT Sham (0.383 ± 0.225, n = 12) in the MG [p = 0.0161]. rs6265 allele carriers showed no such effect, as rs6265 Estim (0.219 ± 0.076, n = 11) was similar to rs6265 Sham (0.371 ± 0.213, n = 10) [p = 0.0934]. Additionally, WT Estim showed higher amplitudes than both rs6265 Sham [p = 0.0153] and rs6265 Estim [p < 0.0001]. In the LG muscle, WT Estim (0.674 ± 0.316, n = 12) rats showed significantly higher CMAP responses compared to WT Sham (0.318 ± 0.127, n = 12) [p < 0.0001]. In contrast, the averages of rs6265 Estim (0.352 ± 0.109, n = 11) and rs6265 Sham (0.337 ± 0.123, n = 10) were not significantly different [p = 0.8606]. In the LG, WT Estim animals were also more recovered than both rs6265 Sham [p = 0.0002] and rs6265 Estim [p = 0.0003]. These results indicate that electrical stimulation treatment enhances functional sciatic nerve muscle reinnervation in WT but not rs6265 rats (Figure 2B, Figure 2C).

**Figure 2:**
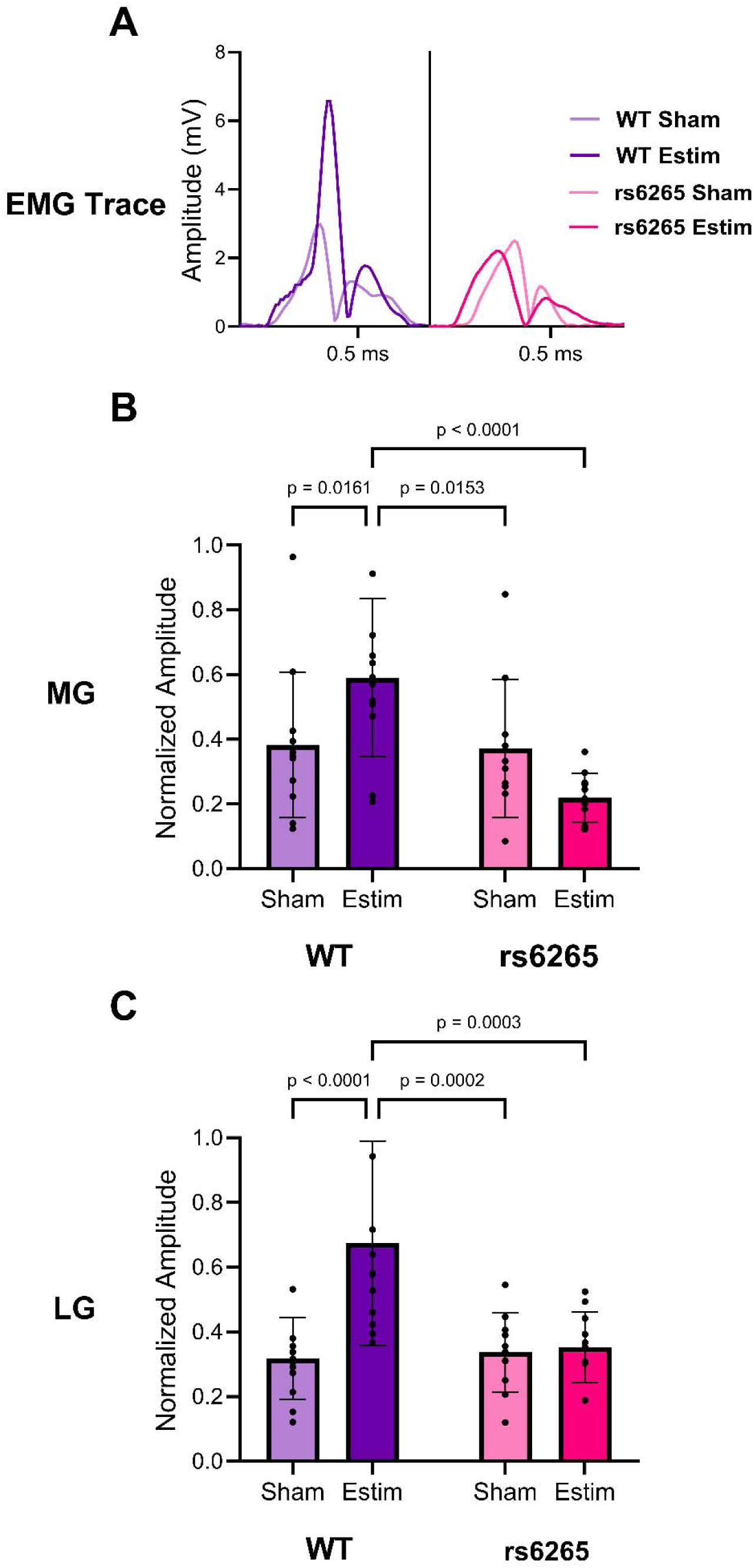
Electrophysiological Analysis of Muscle Reinnervation. **A.** Rectified electromyographic (EMG) trace of lateral gastrocnemius (LG) compound muscle action potential (CMAP) acquired in response to direct stimulation of sciatic nerve eight weeks after injury. Examples are shown for wild-type (WT; left) and rs6265 (right) allele carriers with electrical stimulation (Estim) and Sham treatments. **B.** CMAP amplitude of medial gastrocnemius muscle (MG) eight weeks post-injury normalized to the baseline value. **C.** Normalized CMAP amplitude of LG at eight-week time point.

### Muscle Fiber Analysis: Electrical Stimulation Treatment Leads to Greater Fiber Cross-Sectional Area in Wild-Type but not rs6265 Allele Carriers

MGs were sectioned, stained, and imaged to evaluate the average FCSA. Example images are shown in Figure 3B. Additionally, 12 uninjured rat MGs (six WT and six rs6265 allele carriers, three male and three female in each) were sectioned, stained, and analyzed as a comparison.

**Figure 3:**
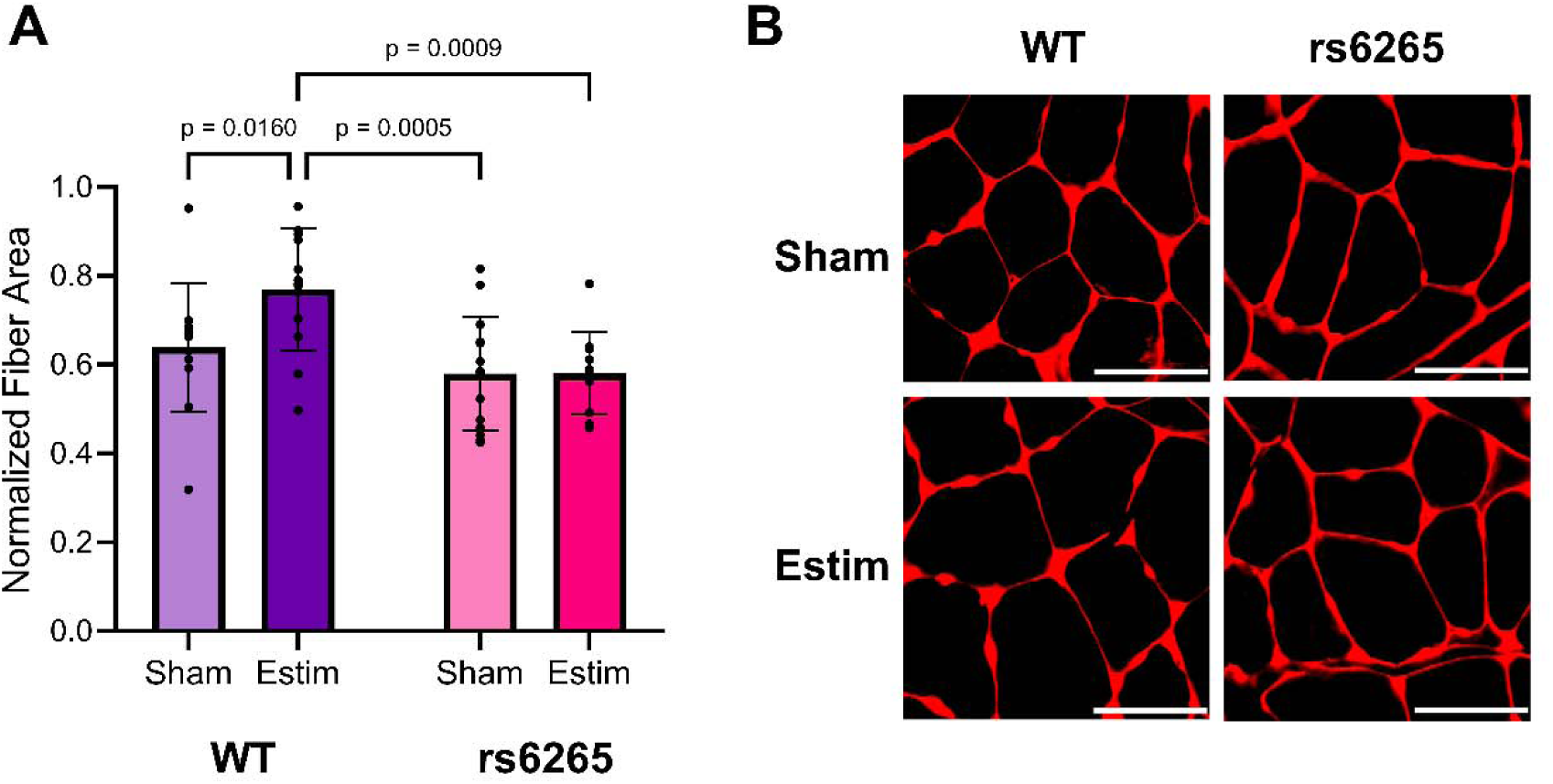
Muscle Fiber Cross-Sectional Area Analysis. **A.** Eight-week muscle fiber cross-sectional area of medial gastrocnemius muscle (MG). Average fiber areas are normalized to genotype- and sex-matched uninjured controls. **B.** 40x images of muscle fibers from MG cross-sections stained with Collagen IV (red). Scale bar = 50 µm.

Uninjured WT males had an average FCSA of 978 ± 34.2 μm^2^, while WT females had 739 ± 152 μm^2^. Uninjured rs6265 males had muscle fibers sized 1082 ± 131 μm^2^, and rs6265 females had 768 ± 47.0 μm^2^. Due to sex-dependent differences in fiber size, FCSA values for injured rats were normalized to uninjured animals of the same sex and genotype. These normalized values were evaluated for effect of sex using a three-way ANOVA, which was not found to be significant [p = 0.4645], so the data for both sexes were combined. Using a two-way ANOVA, WT Estim rats (0.770 ± 0.138, n = 12) were found to have larger muscle fiber areas than WT Sham rats (0.639 ± 0.145, n = 12). rs6265 carriers did not show an effect of treatment, as the rs6265 Estim (0.581 ± 0.092, n = 11) and rs6265 Sham (0.580 ± 0.128, n = 14) animals had similar values [p = 0.9815]. Additionally, the WT Estim group had significantly greater muscle fiber area than both rs6265 Sham [p = 0.0005] and rs6265 Estim [p = 0.0009]. Thus, TES led to greater FCSA in WT but not rs6265 allele carriers (Figure 3A).

### Bursting Frequency and Weighted Firing Rate: iPSC Motor Neurons Generated from Both Lines Develop Similar Levels of Activity over 25 Days Post-Seeding

Following motor neuron differentiation, we examined baseline levels of weighted firing rate (Figure 4E) and bursting frequency (Figure 4F) for both lines over 25 days post-seeding. Data are shown for every five days, starting at day five. The WT (Val/Val) line had a weighted firing rate of 0.447 ± 0.444 (n = 15) at day 5, 0.642 ± 0.537 (n = 47) at day 10, 0.875 ± 0.768 (n = 55) at day 15, 1.251 ± 1.042 (n = 65) at day 20, and 1.465 ± 1.477 (n = 61) at day 25. The rs6265 (Val/Met) line had a weighted firing rate was 0.301 ± 0.281 (n = 10) at day 5, 0.918 ± 0.501 (n = 25) at day 10, 1.156 ± 1.000 (n = 35) at day 15, 1.576 ± 1.177 (n = 38) at day 20, and 1.540 ± 1.267 (n = 39) at day 25. Weighted firing rate significantly increased over time [p < 0.0001] but did not differ between the lines [p = 0.1883]. WT bursting frequency at day 5 was 0.008 ± 0.000 (n = 2), at day 10 was 0.081 ± 0.084 (n = 14), at day 15 was 0.108 ± 0.087 (n = 22), at day 20 was 0.092 ± 0.077 (n = 31), and at day 25 was 0.107 ± 0.134 (n = 29). rs6265 bursting frequency at day 5 was 0.013 ± 0.006 (n = 2), at day 10 was 0.104 ± 0.116 (n = 13), at day 15 was 0.101 ± 0.112 (n = 19), at day 20 was 0.135 ± 0.112 (n = 29), and at day 25 was 0.126 ± 0.121 (n = 30). A two-way ANOVA found that there was no difference in bursting frequency over time [p = 0.3581] or between genotypes [p = 0.5218]. This data shows that the baseline activity level of both strains is similar and allows use of this model to study activity-dependent BDNF release between genotypes.

**Figure 4:**
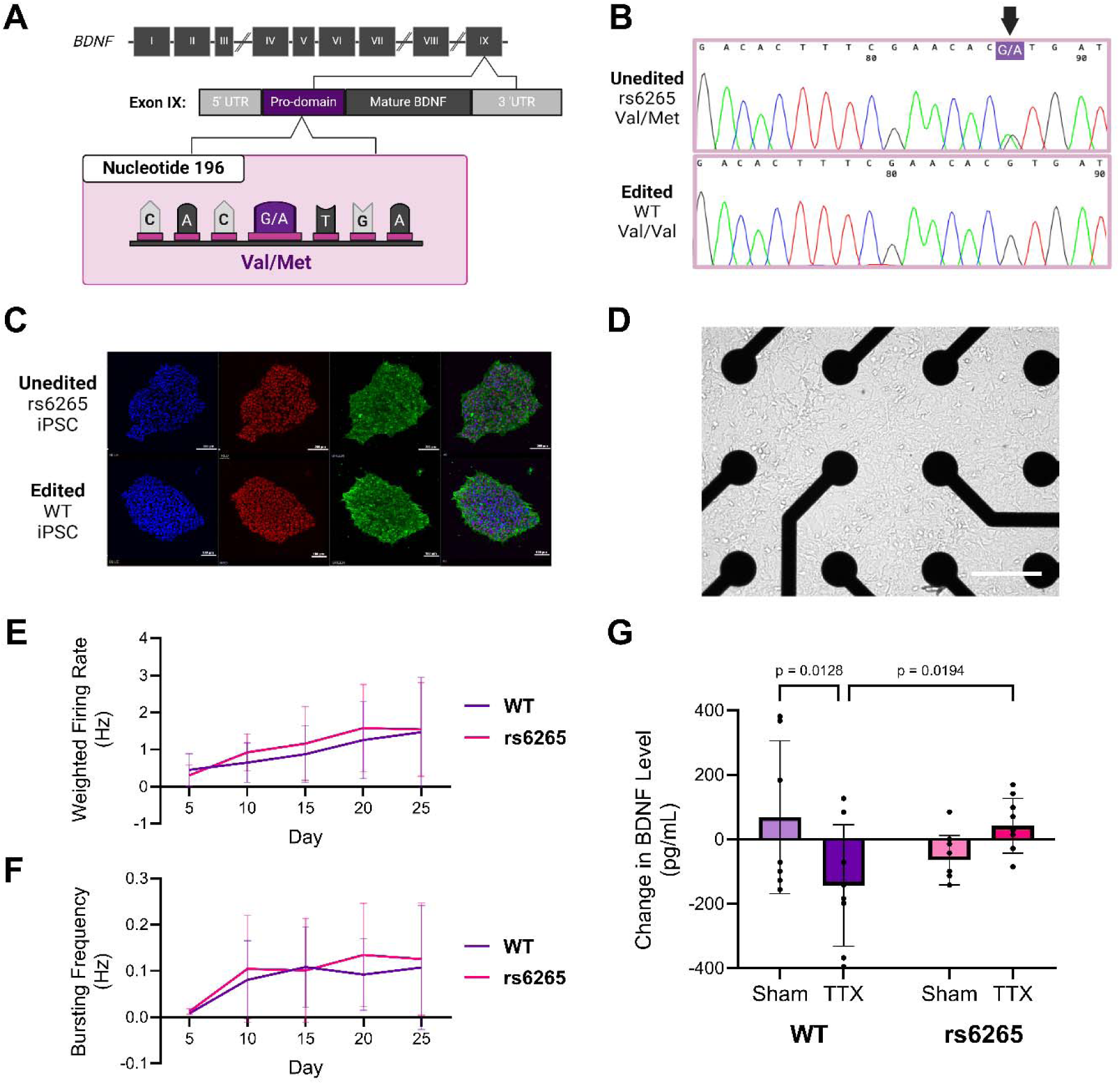
Evaluation of BDNF Secretion in WT and rs6265 Motor Neurons. **A.** A single nucleotide polymorphism (SNP) in the terminal exon of the brain-derived neurotrophic factor (BDNF) gene, shown here in purple, dictates the appearance of the wild-type (WT) and rs6265 alleles. Created with BioRender.com. **B.** Chromatograms confirming CRISP/Cas9 gene editing of a human induced pluripotent stem cell (iPSC) line that carries the rs6265 (Val/Met, G/A) genotype is changed to WT (Val/Val, G/G), black arrowhead. **C.** Reprogrammed iPSCs with either the rs6265 or WT genotype stained for DAPI in blue, SSEA4 in green, and OCT4 in red to show pluripotency prior to differentiation. **D.** Motor neurons are differentiated from isogenic iPSC lines and seeded on multielectrode arrays. Scale bar = 100 µm. **E.** Weighted firing rate and **F.** bursting activity, showing that iPSC motor neurons from both lines develop similar levels of spontaneous electrical activity over 25 days post-seeding. **G.** Difference in BDNF secretion level at baseline and 24 hours after tetrodotoxin (TTX) application to block electrical activity.

### rs6265 Polymorphism Impairs Activity-Dependent BDNF Release in WT Allele Carriers: TTX Application Blocks BDNF Secretion in WT Allele but Not in rs6265 Allele Line

We studied BDNF release in WT allele and rs6265 allele cell lines to evaluate gene-dependent secretion. Since BDNF is released by both activity-dependent and activity-independent (constitutive) mechanisms, the decrease in BDNF levels following activity suppression by TTX provides insight into the activity-dependent pathway, which is known to be compromised in rs6265. TTX was applied to both WT and rs6265 lines to block electrical activity. Since secreted BDNF is relatively unstable in culture, the change in BDNF level (pg/mL) between baseline and 24 hours post-treatment mainly reflects the suppression of its activity-dependent secretion. The WT allele shows a significant reduction in BDNF following TTX application (−143.163 ± 188.711, n = 8) when compared to the Sham (68.678 ± 236.941, n = 7) [p = 0.0128]. In contrast, the rs6265 allele in the TTX (42.419 ± 84.542, n = 9) and Sham (−64.413 ± 76.833, n = 9) condition have similar values [p = 0.1527] (Figure 4G). Additionally, the rs6265 allele post-TTX value is significantly greater than the WT allele post-TTX [p = 0.0194]. The WT line has reduced BDNF after its activity is suppressed with TTX treatment, but the rs6265 line has no change in activity following the same treatment. These results are consistent with the hypothesis that the rs6265 SNP impairs activity-dependent BDNF release.

## Discussion

Our study presents evidence on a gene-treatment interaction between the BDNF rs6265 genetic polymorphism and the efficacy of TES, a therapy known to enhance peripheral axon regeneration in humans and preclinical models^25^. This has notable clinical implications since this SNP that reduces BDNF, essential for the survival and grown of neurons in development^26,27^, is carried by nearly a third of the world population^14^. rs6265 allele carriers exhibit impaired activity-dependent secretion of BDNF, which is a mechanism that has been associated with TES effect on axon regeneration^18,19^. Consistent with this, our study finds that unlike WT allele motor neurons in vitro, BDNF secretion of rs6265 carriers is not reduced when their spontaneous firing activity is blocked using a voltage-gated sodium channel blocker such as TTX. Furthermore, our in vivo results show that the presence of the rs6265 allele attenuates the beneficial effects of TES in a rat model of muscle reinnervation after PNI. Taken together, these findings support the concept that patient specific factors such as genotype, should be prioritized for more comprehensive examination with respect to clinical outcomes after PNI and response to new and emerging treatments.

Our findings align with previous preclinical research that indicate carriers of the rs6265 allele exhibit a differential response to activity-dependent treatments after PNI. McGregor et al.^28^ investigated the efficacy of therapeutic exercise in a mouse model of peripheral axon regeneration with rs6265 transgenic mice. Their study demonstrated that SNP allele carriers do not benefit from an activity-dependent treatment, treadmill exercise, which has been previously shown to augment PNI outcomes in rodents^29–32^. Similar to our study, McGregor et al. used an in vitro model to demonstrate the impaired activity-dependent secretion of BDNF of rs6265 carriers and showed this by applying optogenetic stimulation of dorsal root ganglia isolated from mice that expressed an excitatory ion channel that was sensitive to blue light^28,33^.

However, one notable difference is that McGregor et al. found that under control conditions (without TES) the rs6265 mice exhibited earlier reinnervation of the LG compared to WT mice, which we did not observe in our rat model. This observation, however, is aligned with recent data from our group demonstrating a paradoxical enhancement of neurite outgrowth when WT embryonic dopamine neurons were engrafted into parkinsonian rs6265 rats compared to WT neurons engrafted into WT graft recipient rats^21^. These findings suggest a beneficial role for the variant rs6265 allele in axonal growth and regeneration such that even WT neurons can be stimulated to show enhanced neurite outgrowth when transplanted ectopically into an rs6265 allele-carrying host. It is of note that the valine to methionine substitution associated with the rs6265 SNP lies within the pro-domain of BDNF, and when pro-BDNF is cleaved, active mature BDNF and the pro-peptide are liberated. While little is known about the BDNF pro-peptide, it has been suggested to act as a novel biomolecule, which could be responsible for the beneficial role of the variant rs6265 allele in axonal growth and regeneration^34^.

The reason for the difference between our study and McGregor et al. is not clear but may be due to motor unit subtype specific effects in BDNF. For example, one study showed that the exercise-related increase in BDNF secretion in tibialis anterior and MG muscles of wild-type rats is not seen in the soleus muscle^35^. The rat soleus muscle is almost exclusively innervated by slow type motor units^36^, whereas mice have a much higher proportion of fast motor unit subtypes in the lower leg muscles^37^. This difference in motor unit composition may underlie the varying responsiveness to BDNF, potentially contributing to the observed discrepancies in regenerative outcomes across studies, although this remains speculative. Cumulatively, these observations suggest that axon regenerative capacity may reflect cell intrinsic properties of different motor neuron subpopulations as well as non-cell autonomous events regulated by surrounding cell types and the global environment.

Although genetic interactions with human PNI outcomes are not well-studied, the rs6265 SNP has also been implicated in modifying outcomes of other types of neurotrauma. Studies have shown that the rs6265 allele can be protective in patients with severe traumatic brain injury (TBI). For example, male Vietnam War veterans with penetrating TBI who carried the rs6265 allele exhibited better executive function, general intelligence, working memory, and processing speed compared to WT (Val/Val) individuals in long-term follow-up after the injury^38^. It is hypothesized that this benefit may arise from the reduced secretion of pro-BDNF in rs6265 carriers, which may mitigate the apoptotic effects of pro-BDNF. Pro-BDNF preferentially binds to the p75 receptor, which has pro-apoptotic and synapse-eliminating effects, rather than the canonical BDNF receptor TrkB, which promotes neuronal survival and growth^39^. However, the interaction between the rs6265 SNP and neurotraumatic injuries is certain to be more complex. In a separate study of severe TBI patients, the rs6265 allele, along with another high-risk BDNF polymorphism (rs7124442), initially reduced the risk of acute death following injury (within seven days). At longer time points (up to 365 days), the risk of death in rs6265 carriers differed by age. Specifically, carriers of both the rs6265 allele and rs7124442 SNP with severe TBI had relatively reduced death rates if they were over 45 years of age, but relatively increased death rates if they were under 45 years of age^40^. The underlying reasons for these gene-age-trauma interactions remain largely speculative and require further confirmation by larger studies in future.

Despite the results presented here on the interaction between the rs6265 SNP and the efficacy of TES for peripheral axon regeneration, several limitations should be acknowledged. Firstly, our experiments were conducted using a rat model, which, while informative, does not fully replicate the complexities of human physiology and genetic diversity. Additionally, the study focused on a single genetic SNP (rs6265), and the interaction of this SNP with other genetic and environmental factors remains unexplored. Addressing these limitations will be crucial in future research. In the context of clinical applications, our findings underscore the potential need for more personalized treatment strategies. Future clinical trials will likely benefit from incorporating stratification based on patient-specific factors such as genotype. This approach can help identify subgroups of patients who are likely to benefit most from TES and other activity-dependent treatments, thereby optimizing therapeutic outcomes. For instance, genetic screening for the rs6265 SNP could be used to tailor interventions, ensuring that patients with the rs6265 allele receive alternative or adjunctive therapies that compensate for their impaired BDNF secretion.

In summary, our study demonstrates that the rs6265 SNP significantly influences the efficacy of TES in promoting peripheral axon regeneration. These findings highlight the importance of considering patient-specific genetic factors in both preclinical research and clinical trial design. By adopting a personalized approach to treatment, we can enhance recovery outcomes and reduce variability in response to therapies for peripheral nerve injuries.

## Acknowledgements

Alyssa Weston, MSc for her technical assistance with stem cell cultures and differentiations.

## Ethical Publication Statement

We confirm that we have read the Journal’s position on issues involved in ethical publication and affirm that this report is consistent with those guidelines.

## Abbreviations

BDNF: brain-derived neurotrophic factor
CMAP: compound muscle action potential
CSA: cross-sectional area
EMG: electromyography
Estim: electrical stimulation
FCSA: fiber cross-sectional area
iPSC: induced pluripotent stem cell
LG: lateral gastrocnemius muscle
MG: medial gastrocnemius muscle
NGS: next generation sequencing
PDL: poly-D-lysine
PNI: peripheral nerve injury
SNP: single nucleotide polymorphism
ssODN: single-stranded oligodeoxynucleotide
TBI: traumatic brain injury
TES: therapeutic electrical stimulation
TIB: tibia
TTX: tetrodotoxin
WT: wild-type

